# Changes in spectral signature of leaves after desiccation: Implications for the prediction of leaf traits and plant-soil interaction in herbarium samples

**DOI:** 10.1101/2024.01.21.576284

**Authors:** Natalia L. Quinteros Casaverde, Shawn P. Serbin, Douglas C. Daly

## Abstract

- The study investigated the impact of specimen desiccation on spectral signatures of plant tissue and its influence on models predicting biochemical and physiological traits, as well as ecosystem function.
- Analyzing 56 species, this study quantified the impact of leaf desiccation on: (1) the degree of change in the reflectance intensity, and its first and second derivatives within the visible - shortwave infrared spectrum, (2) the difference in change between wavebands used by Radiative Transfer Models (RTMs) for predicting traits in fresh leaves, (3) the prediction of leaf traits using PROSPECT RTM, and (4) the prediction of soil nutrient components from the leaf.
- Comparing desiccated specimens to fresh leaves, this study found the highest degree of change within the near infrared for the reflectance intensity, its first and second derivatives. Specific wavebands used for the prediction of leaf traits changed less that others in the VIS-SWIR. The prediction uncertainty for PROSPECT RTM differed for various leaf traits, showing an increase for equivalent water thickness, and carotene content, and a decrease in brown pigments and dry mass. Leaf traits predicted from desiccated leaves were better at predicting relationships with soil components, such as soil chemical properties and macro and micronutrients than fresh leaves.
- Desiccated leaves preserve and improves the capacity to inform us about important aspects of ecosystem functioning, particularly nutrient transfer between leaf and soil.

## Introduction

Studies of ecosystem functioning have grown substantially in recent years, at least partially due to the availability of novel ways to collect critical plant functional trait data (Durán et al., 2019; Jetz et al., 2016; Martin et al., 2017; Ustin & Gamon, 2010). One such advancement is the use of leaf reflectance spectroscopy to predict functional traits from fresh and desiccated leaf spectral reflectance (Ely et al., 2019; Kothari et al., 2023; Schweiger et al., 2018; Serbin et al., 2014; Z. Wang et al., 2020). *In situ* measurements are constrained by time or other resources, but spectroscopic techniques can provide rapid, cost-efficient estimates of functional traits, even after considering the price of a spectrometer, the costs associated with laboratory equipment, time, and resources required for wet chemical and analytical techniques (Costa et al., 2018; Kothari et al., 2023).

While these methods are useful when applied to freshly collected leaves, the changes in leaf reflectance patterns expected after leaf clipping are poorly known. While leaf aging and phenological fluctuations are relatively well comprehended (Foley et al., 2006), leaf reflectance patterns are not entirely understood for desiccated leaves. As such, the impact of leaf desiccation on trait predictability using biophysical models is poorly understood. Fabre et al. (2011) showed that water disrupts solar radiation within the leaf across the visible and shortwave-infrared continuum. Therefore, understanding the spectral changes after leaf desiccation, and the consequences for trait prediction, are of interest to the scientific community due to the possible impact on ecosystem function inference (Kothari & Schweiger, 2022).

Because spectroscopic approaches can be non-destructive, they can provide a method that is easily applicable to samples kept in herbaria, which around the world safeguard priceless plant collections since before the industrial revolution (Campos et al., 2021; Nualart et al., 2017). In addition to having information about the plant identity, leaves carry the footprint of species habitats and their responses to changing environments in their external and internal structures (Oliveras et al., 2020; Violle et al., 2007). When a leaf is collected and desiccated for an herbarium sample, a snapshot of the time and the place it lived in is preserved (Willis et al., 2017). But leaf desiccation involves eliminating water through multiple means (Freschet et al., 2010): leaves are pressed and then exposed to a heat source, causing the leaf to transpire while within the plant press, until it is dry (Blanco et al., 2006).

The literature on leaf spectroscopy suggests that desiccated leaves still contain enough spectral information to provide insight about their taxonomic identity (Lang et al., 2015, 2017). and, potentially, to enable leaf trait prediction (Kothari et al., 2023; Z. Wang et al., 2020). Lang et al. (2015, 2017) showed that it is possible to classify plants to different taxonomic levels based on the near-infrared spectrum. There is also evidence that nitrogen content, related to enzymes and photosynthetic pigment, is better predicted with the spectral reflectance of desiccated rather than fresh leaves (Kothari et al., 2023; Z. Wang et al., 2020). Similarly, the prediction of leaf dry mass, carbon, and leaf nutrients improves after leaves are desiccated (Kothari et al., 2022). In contrast, nothing is known about the impact of leaf desiccation on biophysical models that infer traits from leaf spectra, such as leaf-level radiative transfer models. This is in stark contrast with trait prediction based on the spectral reflectance of fresh leaves: Wang et al. (2023) found that both statistical and radiative transfer models predicted leaf traits with high accuracy based on the spectral reflectance of fresh leaves. Therefore, an evaluation of biophysical model performance in desiccated leaves is needed to continue advancing the field.

Different or fluctuating environmental conditions leave a trace within the spectral signature of leaves (400-2400 nm) in areas of maximum absorption of photosynthetic pigments (at 430 and 660 nm), water (at 1450 and 1980 nm), protein (1980 and 2350 nm), and cellulose (at 2350 nm). These areas are coined as local maxima by Schweiger et al. (2018), due to their high variability when plants are exposed to different environmental conditions. These local maxima are used in biophysical models to simulate leaf spectral reflectance, and in consequence, its inversion lets us predict back these traits (Jacquemoud et al., 1996). Within leaves, water strongly influences the behavior of the radiant energy in the short-wave infrared spectrum, particularly at 1450 nm and 1940 nm, as well as the behavior of biophysical models such as radiative transfer models (RTM) (Zarco-Tejada et al., 2003). Thus, it is reasonable to hypothesize that these local maxima will be located at the same wavelengths in desiccated leaves. Leaf desiccation not only eliminates water from within the leaf, but it also initiates the degradation of photosynthetic pigments that absorb solar radiation within the visible (VIS) spectrum (Peñuelas & Inoue, 1999), proteins, and other cell components with the subsequent development of brown pigments, such as phenolic compounds, which absorb radiant energy in the infrared spectrum (Féret et al. 2008). The increase of these brown pigments is sensed in the near infrared (NIR) below 1300 nm (Féret et al., 2008). Another factor associated with increased NIR reflectance is the reduction of leaf thickness due to drying cells’ contraction (Foley et al., 2006). Therefore, WEcan expect that when components of the leaf vary, for instance through desiccation, the absorption features will change, affecting the shape (e.g., slope and curvature) and the intensity of the spectral signatures, the wavelength regions showing maximum variation (Jacquemoud & Ustin, 2019), as well as the precision of radiative transfer models that predict biochemical and structural traits (Jacquemoud et al., 1996).

Investigating the utility of these trait-predicting radiative transfer models in desiccated leaves can allow us to substantially improve our understanding of how ecosystems function. Critical to ecosystem functions are belowground processes such as decomposition, soil carbon storage, and nutrient cycling, which have proven to be elusive targets for detection (Cavender-Bares, Schneider, et al., 2022; Cavender-Bares, Schweiger, et al., 2022). Since the internal changes in the leaf are the consequence of the interaction between plants and their environment (above and belowground), the spectral reflectance of leaves can provide useful information about those processes (Kattenborn et al., 2017, 2019; Kattenborn & Schmidtlein, 2019). The replenishment of nutrients is fundamental to balance the content of leaf functional traits and plants interact with the soil most intensely in the rhizosphere to acquire essential nutrients (Liu et al., 2022). Studies about plant-soil interactions generally focus on the impact of soils on plant traits through the manipulation of soil nutrients (Ehrmann & Ritz, 2013; Ordoñez et al., 2009), but spectroscopy of leaves can help us elucidate aspects of the nutrient cycling between leaves and soil, such as nutrient transfer, since leaf trait balance appears to partially drive belowground nutrient transfer along with decomposers (Madritch et al., 2020; Perez-Moreno & Read, 2001; Tedersoo & Bahram, 2019; Zhao et al., 2021). Therefore, as the portion of solar radiation that leaves reflect into the environment correlates with leaf traits influenced by soil nutrient availability, leaf traits predicted from spectral reflectance can serve as a valuable bridge to connect these ecological interactions. More specifically, plants need macro (nitrogen, magnesium, phosphorus, sulfur, potassium, and calcium) and micronutrients (Iron, copper, zinc, molybdenum, boron, chlorine, and nickel) from the soil for biosynthesis and maintenance of photosynthetic pigments, enzymes, and structural components (Maathuis, 2009). All these nutrients are not freely available to plants in the soil because they are bound to soil particles by their opposite electrical charges (Drake et al., 1951). The process in which these nutrients are released and pumped into the root is through the process of cation exchange (Mattson et al., 1950; Rufyikiri et al., 2007; Williams & Coleman, 1950) or soil acidification (Chaney et al., 1972) imposed by the plants around their root systems guided by a mechanism that allowed the plant to regulate nutrient intake (Amtmann & Rubio, 2012). Therefore, it is reasonable to expect a strong relationship between the spectrally predicted leaf traits and soil elements, allowing for the prediction of soil components through these leaf constituents.

In this investigation we ask (1) whether the wavebands associated with the highest variation of photosynthetic pigments, water, proteins, and dry mass in fresh leaves (a.k.a. local maxima as per Schweiger et al., 2018) are different from the wavebands of highest variation in dry leaves, (2) whether radiative transfer models, and more specifically PROSPECT-D RTM, can predict, in dry leaves, the natural course of the decrease in photosynthetic pigments and water, as well as the increase of leaf dry mass, brown pigments, and the structural parameter, a proxy of leaf internal complexity (Spafford et al., 2021), (3) whether leaf trait predictions from PROSPECT-D RTM will have higher precision for traits whose concentration increase in response to desiccation, and lower precision for leaf traits whose concentration decrease with desiccation, (4) whether leaf traits predicted from dry leaves will be better at capturing the variation of soil properties such as macronutrients (nitrogen, magnesium, phosphorus, sulfur, potassium, and calcium), micronutrients (iron, copper, zinc, and boron), aluminum, and chemical characteristics of the soil such as the cation exchange capacity and the pH within the rhizosphere relative to freshly collected leaves.

## Methods

To evaluate the consequences of leaf desiccation on leaf trait predictability through PROSPECT-D RTM, and to estimate the co-variation between soil nutrients and leaf traits, we collected three sun-lit leaves from individual trees belonging to 56 species (Table 1) growing in the Enid A. Haupt Conservatory at the New York Botanical Garden (NYBG). To measure the spectral reflectance of the leaves while still fresh, after clipping them from the tree, we used a Spectra Vista Corporation (SVC HR 1024i) spectroradiometer connected to an SVC LC-RP-Pro leaf clip with an internal full-spectrum light source, calibrated using a Spectralon® white reference. To obtain the spectral reflectance of desiccated leaves, we first dried them by carefully placing each one within a 30.5 x 46 cm plant press, positioned between C-flute corrugated ventilators, and exposed to a small room heater with the temperature set at its maximum and for 72 hours (Figure S1), following Blanco et al. (2006). Leaves placed in the middle of the plant press were moved to the ends every 12 hours to allow for an even desiccation process. Measurements of the spectral reflectance of the desiccated leaves were then obtained by following the same protocol used when they were fresh.

**Table 1.** Sampled species organized by family and species name.

| Family | Scientific Name (Code) |
| --- | --- |
| ACANTHACEAE | *Aphelandra flava* Nees, *Aphelandra tetragona* (Vahl) Nees, *Louteridium donnell-smithii* S. Watson, *Megaskepasma erythrochlamys* Lindau, *Pachystachys coccinea* (Aubl.) Nees, *Ruellia speciosa* Mart. ex Nees, *Sanchezia sanmartinensis* Leonard & L.B. Sm., *Sanchezia speciosa* Leonard, *Thunbergia grandiflora* Roxb., *Thunbergia mysorensis* (Wright) T. Anderson |
| ERICACEAE | *Anthopterus cuneatus* A.C. Sm., *Cavendishia lebronae* Luteyn, *Diogenesia amplexans* (Sleumer) Sleumer, *Macleania bullata* Yeo, *Macleania coccoloboides* A.C. Sm., *Macleania pentaptera* Hoerold, *Macleania rupestris* (Kunth) A.C.Sm., *Psammisia ramiflora* Klotzsch |
| FABACEAE | *Brownea macrophylla* hort. ex Mast., *Calliandra haematocephala* Hassk., *Calliandria tergemina* (L.) Benth., *Hymenaea* spp. (HYspp), *Inga edulis* Mart., *Mucuna bennettii* F. Muell., *Pterocarpus officinalis* Jacq., *Samanea tubulosa* (Benth.) Barneby & J.W. Grimes, *Strongylodon macrobotrys* A. Gray, *Tamarindus indica* L., *Tipuana tipu* Kuntze |
| MYRTACEAE | *Campomanesia guazumifolia* (Cambess.) O. Berg, *Eugenia uniflora* L., *Pimenta racemosa* (Mill.) J.W. Moore, *Psidium friedrichsthalianum* (O. Berg) Nied., *Psidium guajava* L., *Psidium guineense* Sw., *Psidium cattleianum* Sabine |
| RUBIACEAE | *Augusta rivalis* (Benth.) J.H. Kirkbr., *Cinchona* spp., *Coffea arabica* L., *Coffea canephora* Pierre ex A. Froehner, *Deppea splendens* Breedlove & Lorence, *Exostema nitens* Urb., *Genipa americana* L., *Hamelia cuprea* Griseb., *Osa pulchra* (D.R. Simpson) Aiello, *Psychotria nervosa* Sw., *Posoqueria latifolia* (Rudge) Roem. & Schult., *Pogonopus speciosus* (Jacq.) K. Schum., *Rosenbergiodendron longiflorum* (Ruiz & Pav.) |
| SOLANACEAE | *Brunfelsia grandiflora* D.Don, *Brunfelsia nitida* Benth., *Brunfelsia undulata* Sw., *Cestrum aurantiacum* Lindl., *Iochroma cyaneum* (Lindl.) M.L. Green ex G.H.M. Lawr. & J.M. Tucker, *Iochroma fuchsioides* (Bonpl.) Miers, *Solandra* spp. |

To understand the changes in the leaf spectral signature caused by desiccation, we calculated the first and second derivatives of the reflectance as a measure of the slope of the tangents and curvature (Louchard et al., 2002) between two wavebands along the spectral signatures of both fresh and dry leaves. Before computing these metrics, the wavebands between 1895 nm and 1905 nm were removed from the analysis to exclude an unnecessary splice correction performed by the spectral preprocessing pipeline (Frey et al., 2020). To quantify changes in the leaf spectral signature caused by desiccation, we calculated the Manhattan distance as a measure of dissimilarity between individual spectral reflectance, between the first derivatives of the spectral reflectance, and between its second derivatives, comparing samples obtained from fresh and desiccated leaves. For that we used the dist2 function from the R package flexclust (Leisch, 2006), and used the diagonal of the distance matrices to find the bands with the highest degree of change. The formula used to obtain the Manhattan distances was the sum of the absolute difference between two samples at each waveband. This metric accommodates the high degree of autocorrelation in spectral data (Schweiger et al., 2018). Then, to test whether the range of dissimilarity in wavebands presenting the highest degree of change after leaf desiccation was statistically different from the range of dissimilarity in wavebands known to be absorbed by important leaf components (such as photosynthetic pigments, water, proteins, and cellulose), WE ran a bootstrap permutation to obtain the error and 95% confidence intervals around the dissimilarity values at those target wavelengths (2.5th and 975th percentiles) with the boot and boot.ci functions from the R package boot (Ripley 2022), and evaluated the confidence intervals. To test whether wavebands with overlapping confidence intervals were statistically different, we ran a bootstrap t-test with the function boot.t.test from the R package rfast (Tsagris & Papadakis, 2018). Finally, the values of the diagonals from the reflectance, the first derivative, and the second derivative were normalized and compared.

To test for changes in precision of the trait prediction in PROSPECT-D RTM after leaves were desiccated, we calculated the following leaf parameters: internal complexity (structural N), total chlorophyll content (Cab g/m^2^), carotene content (Car g/m^2^), dry matter content (Cm g/m^2^), water content (Cw g/m^2^), and brown pigment content (Cbrown g/m^2^), following Shiklomanov et al. (2016) pipeline, for both fresh and desiccated leaves. We trained the model with libraries obtained from four leaf optical property experimental databases: LOPEX (Hosgood et al., 1993), ANGERS (Jacquemound et al., 2003), HAWAII, and CALMIT (Féret et al., 2008). We used prediction uncertainty to evaluate the precision of each trait prediction: this is a metric for error, calculated as the width of the 95% confidence interval relative to the mean value obtained from PROSPECT, through a Bayesian inversion (Shiklomanov et al., 2016). To test whether the uncertainties changed and whether the estimated leaf traits followed natural decay expectations after the leaves were desiccated, we used a paired t-test, using the function paired from the R package PairedData.

Finally, to test whether we can predict the variation of soil nutrients based on traits estimated from the spectral reflectance of fresh and dry leaves, we collected soil samples from the first 10 cm of the rhizosphere of eight individuals of different species: *Osa pulchra* (D.R. Simpson) Aiello (Rubiaceae), *Pimenta racemosa* (Mill.) J.W. Moore (Myrtaceae), *Psychotria nervosa* Sw. (Rubiaceae), *Brunfelsia grandiflora* D. Don (Solanaceae), *Cinchona spp* (Rubiaceae), *Eugenia uniflora* L. (Myrtaceae), *Brownea macrophylla* hort. ex Mast. (Fabaceae), *Aphelandra tetragona* (Vahl) Nees (Acanthaceae), with each species growing in a different soil bed. For that, we used a soil core sampler with a diameter of 3 cm and stored in a Nasco Whirl-pak bag (Figure S2). The University of Connecticut Soils Lab conducted soil tests to quantify macro (Nitrate, S, P, K, Ca, and Mg) and micronutrients (B, Cu, Fe, Mn, and Zn), aluminum, soil pH, and cation exchange capacity (CCC) for these eight samples. All soil macro and micronutrients, except nitrate, were obtained through the modified Morgan procedure (McIntosh, 1969), nitrate content was estimated trough the cadmium reduction procedure (Cortas & Wakid, 1990), CCC was estimated through the Ammonium Acetate method at neutral pH (Chapman, 2016), and pH inference through an electronic pH meter (Mclean, 2015). We then built bivariate linear models for fresh and dry leaves separately, evaluating how well the leaf trait predicted soil nutrients and soil properties. For that, leaf trait values were averaged by individual, and coupled with their respective soil nutrient values. The linear model was run with the base R function lm.

## Results

After the leaves were desiccated, the most pronounced changes in reflectance intensity (RI) were detected in the shortwave-infrared ranges of the electromagnetic spectrum. Highest dissimilarity (Figure 1A) was observed at 1894 nm, with a dissimilarity between fresh and dry leaves (Mdist) of 6348.4 (SEboot = 89.2; 95%Ciboot = [6177, 6524]). This was followed by another region of high dissimilarity, observed at 1418 nm, with a Mdist of 5857.03 (SEboot = 128.1; 95%CIboot = [5615, 6118]). The most pronounced changes in the first derivative of the reflectance (D1) were detected in the visible and the shortwave-infrared. Highest dissimilarity between dried and fresh leaves (Figure 1B) appeared at 1386 nm, with a dissimilarity of 95.6 (SEboot = 1.2; 95%CIboot = [93.25, 97.83]), followed by areas of moderate dissimilarity at 720 nm (Mdist = 80.7; SEboot = 2.6; 95%CIboot = [75.69, 85.87]), 1393 nm (Mdist = 94.38; SEboot = 1.1; 95%CIboot = [92.26, 96.56]), and 1874 nm (Mdist = 78.8; SEboot = 1.6; 95%CIboot = [75.64, 81.92]). Desiccation-related changes in the second derivative (D2) of the reflectance were more pronounced within the visible and shortwave-infrared spectrum. Highest dissimilarity for D2 (Figure 1C) was observed at 1888 nm, with a Mdist of 17.7 (SEboot = 0.21; 95%CIboot = [17.28, 18.11]), followed by changes at 402 nm (Mdist = 16.5; SEboot = 1.3; 95%CIboot = [13.92, 19.15]), 728 nm (Mdist = 14.1; SEboot = 0.3; 95%CIboot = [13.6, 14.61]), 1378 nm (Mdist = 14.1; SEboot = 0.2; 95%CIboot = [13.9, 19.15]), 1400 nm (Mdist = 17.7; SEboot = 0.21; 95%CIboot = [17.28, 18.11]).

**Figure 1.**
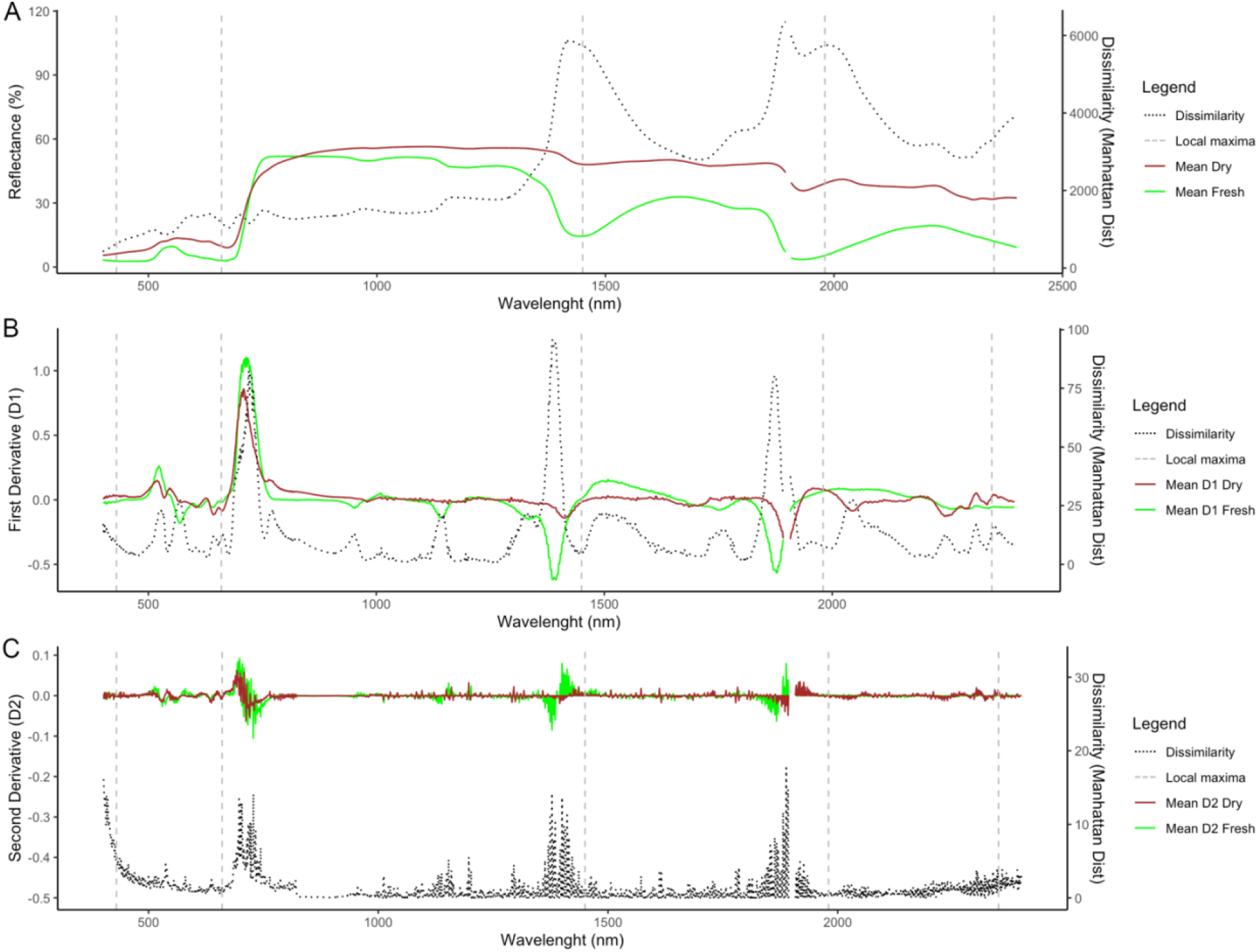
Differences in (A) spectral reflectance intensity, (B) first derivative (slope tangent) of the reflectance, and (C) second derivative (curvature) of the reflectance after leaf dehydration. Grey dashed lines indicate local maxima, *sensu* Schweiger et al. (2018), for fresh leaves, at 430 nm, 660 nm, 1450 nm, 1980 nm, and 2350 nm.

The wavebands of the known local maxima for fresh leaves, as per Schweiger et al. (2018), exhibited lower dissimilarity for the three metrics (RI, D1, and D2) after the leaves underwent the desiccation process (Figure 2 and Figure 3). Bootstrap permutations confirm that the magnitude of those differences are statistically significant, given the lack of overlap in bootstrap confidence intervals, as illustrated in Figure 3. Although the confidence interval of the reflectance intensity at the local maxima for water at 1450 nm overlapped with the highest variability point at 1418 nm (Figure 3A), a paired bootstrap t-test showed that these mean dissimilarities were statistically different (p< 0.05). The evaluation of the response of the spectral metrics to desiccation showed a distinct pattern that emerged after the three metric (RI, D1, and D2) dissimilarities were normalized. The normalized spectral metrics converged to display similar peaks around the wavelength of 1888 nm (Figure 2). These shifts in the spectral signature, indicative of underlying changes in leaf traits, were particularly pronounced in alterations of slope and curvature across the spectrum, as opposed to changes in reflectance intensity. This effect was especially prominent within the visible spectrum (400 to 700 nm), which is sensitive to variations in photosynthetic pigments.

**Figure 2.**
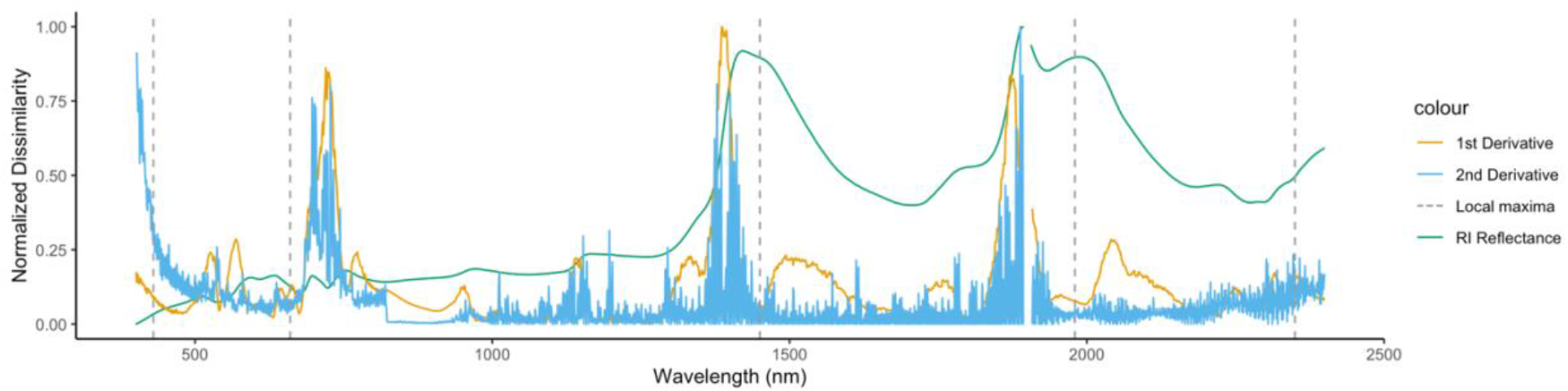
Normalized dissimilarities (Manhattan Mdist) for reflectance intensity, and its first and second derivatives. Grey dashed lines indicate local maxima, sensu Schweiger et al. (2018), at 430 nm, 660 nm, 1450 nm, 1980 nm, and 2350 nm. Grey dashed lines indicate local maxima, *sensu* Schweiger et al. (2018), for fresh leaves, at 430 nm, 660 nm, 1450 nm, 1980 nm, and 2350 nm.

**Figure 3.**
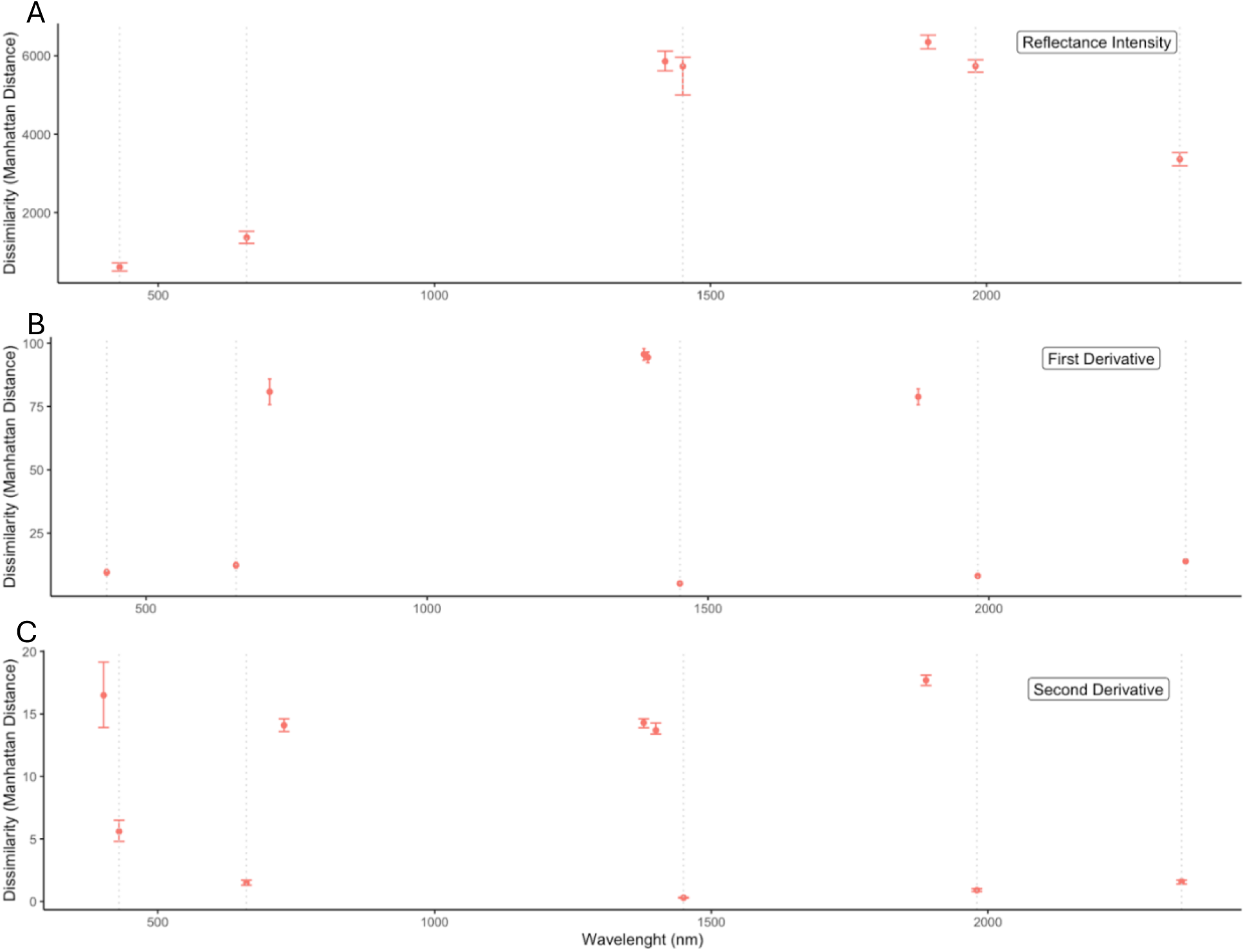
Mean Manhattan distance and bootstrap confidence intervals for bands that presented the highest dissimilarity for (A) the reflectance intensity, (B) first derivative, and (C) second derivative compared to the bands at the local maxima for fresh leaves (as per Schweiger et al., 2018). Dotted lines mark the location for the local maxima for fresh leaves at (430 nm, 660 nm, 1450 nm, 1980 nm, 2350 nm).

After predicting photosynthetic pigment densities, equivalent water thickness, dry mass per area, and brown pigments with PROSPECT-D RTM, the paired t-test on these predicted values, averaged per individual tree, showed that all inferred trait values were statistically different after drying the leaves (p-value < 0.01, Table 2; Figure 4). The same analysis showed an increase in the structural N parameter by 0.64 units, brown pigments by 0.006 g/m^2^, and dry mass by 21.1 g/m^2^. The results for the leaf equivalent water thickness (EWT) and photosynthetic pigments were the opposite; EWT decreased by 198.84 g/m^2^, chlorophyll a & b by 0.18 g/m^2^, and carotene by 0.1 g/m^2^. Correspondingly, the evaluation of the precision of the Bayesian inversion of the PROSPECT-D RTM for trait prediction showed the expected trends. The prediction precision of the estimated trait values in desiccated leaves was higher than the trait values obtained from fresh leaves for structural parameter N, leaf dry mass per area, and brown pigment estimations. In contrast, prediction precision of photosynthetic pigments and water decreased in the desiccated leaves. The paired t-test confirmed that the uncertainty around the prediction of brown pigments and dry matter values decreased in desiccated leaves compared to an increase in carotene and equivalent water thickness uncertainty values (p-value < 0.05, Table 3). But the same test showed that the decrease in uncertainty around the prediction of the structural parameter N and total chlorophyll were not statistically different between fresh and desiccated leaves (Table 3).

**Figure 4.**
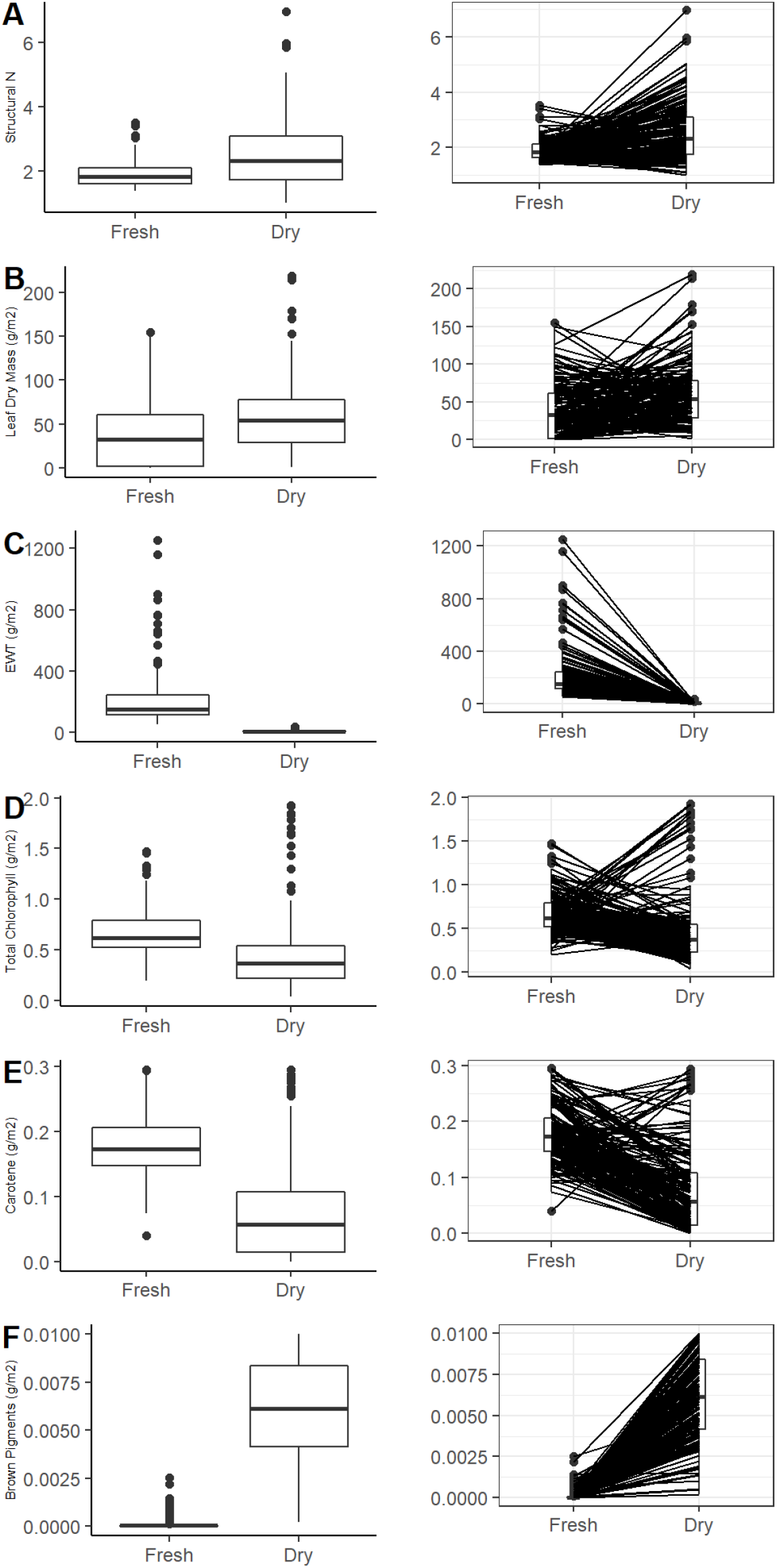
Changes in distribution (right column) and trait value (left column) predicted leaf biochemical and structural traits, before and after desiccation. (A) Structural parameter N, (B) leaf dry mass per area (C) Equivalent Water Thickness (EWT), (D) total chlorophyll content, (E) total carotenoid content, and (F) Brown pigments.

**Table 2.** Evaluation of the trajectory of PROSPECT mean trait values for all individuals after desiccation, based on a paired t-test.

| *Trait* | *Estimate (D)* | *t-stat* | *df* | *LL 95%CI* | *UL 95%CI* | *p-value* |
| --- | --- | --- | --- | --- | --- | --- |
| *Structural N* | 0.64 | 5.3 | 55 | 0.4 | 0.9 | **2.15E-06** |
| *Total chlorophyll (g/m2)* | -0.18 | -3.1 | 55 | -0.3 | -0.07 | **2.9E-03** |
| *Carotene (g/m2)* | -0.1 | -9.3 | 55 | -0.12 | -0.08 | **5.9E-13** |
| *Brown Pigments (g/m2)* | 0.006 | 22.9 | 55 | 0.06 | 0.07 | **2.2E-16** |
| *Equivalent water thickness (g/m2)* | -198.84 | -8.7 | 55 | -244.3 | -153.4 | **5E-16** |
| *Dry mass per area (g/m2)* | 21.1 | 3.24 | 55 | 8.05 | 34.12 | **2E-03** |

**Table 3.** Evaluation of the precision of the Bayesian inversion of PROSPECT RTM based on the mean uncertainty of the prediction for each pair based on paired t-test results.

| *Trait* | *Estimate (D)* | *t-stat* | *df* | *LL 95%CI* | *UL 95%CI* | *p-value* |
| --- | --- | --- | --- | --- | --- | --- |
| *Structural N* | -0.015 | -1.07 | 55 | -0.02 | 0.13 | 2.9E-01 |
| *Total chlorophyll (g/m2)* | 0.06 | 1.42 | 55 | -0.02 | 0.13 | 1.6E-01 |
| *Carotene (g/m2)* | 0.62 | 5.01 | 55 | 0.37 | 0.86 | **5.9E-10** |
| *Brown Pigments (g/m2)* | -3.08 | -18.12 | 55 | -3.41 | -2.74 | **2.20E-16** |
| *Equivalent water thickness (g/m2)* | 0.1 | 3.38 | 55 | 0.04 | 0.15 | **1.43E-03** |
| *Dry mass per area (g/m2)* | -0.77 | -3.31 | 55 | -1.15 | -0.38 | **1.30E-04** |

The outcomes of the linear models analyzing the relationship between soil nutrients and leaf traits predicted from spectral reflectance differed when built with data from fresh vs. dried leaves. Models containing leaf traits predicted from the spectral reflectance of fresh leaves detected a significant relationship between soil manganese and leaf carotene content (p-value < 0.01, R2 = 0.9; Table 2.4), while the remaining leaf traits did not show a statistically significant correlation with soil nutrients or properties (Table A1). Models built with leaf traits predicted from desiccated specimens (Table 4) were nonetheless better at predicting the variation of soil chemical properties and macro and micronutrients. A linear regression showed a statistically significant relationship and a strong correlation between soil pH and leaf brown pigment content (p-value < 0.01, R^2^ = 0.72), as well as between soil phosphorus and leaf carotene content (p-value < 0.01, R^2^ = 0.7). While the relationship between soil sulfur and dry matter content were statistically significant, the correlation was not as strong as the previous soil-leaf relationships (p-value = 0.053, R^2^ = 0.43). Similar results were observed for the relationship between soil boron and brown leaf pigment content (p-value < 0.05, R2 = 0.51), and between soil iron and brown leaf pigment content (p-value < 0.05, R^2^ = 0.43): while the models were statistically significant, the correlations were not as strong.

**Table 4.** List of soil – leaf trait relationships derived from fresh and desiccated specimens which showed statistically significant correlations.

| **Soil Property** | **Soil Trait** | **Leaf Trait<sup>1</sup>** | **Fresh Leaf Model** |  |  | **Dry Leaf Model** |  |  |
| --- | --- | --- | --- | --- | --- | --- | --- | --- |
|  |  |  | **Coefficient** | **R<sup>2</sup>** | **p-value** | **Coefficient** | **R<sup>2</sup>** | **p-value** |
| Chemical property | Soil pH | Cbrown (g/m<sup>2</sup>) |  |  |  | 122.3 | 0.72 | 0.005 |
| Macronutrient | P (g/m<sup>2</sup>) | Car (g/m<sup>2</sup>) |  |  |  | 22.67 | 0.7 | 0.006 |
|  | S (ppm) | Cm (g/m<sup>2</sup>) |  |  |  | -0.19 | 0.43 | 0.047 |
| Micronutrient | Mn (ppm) | Car (g/m<sup>2</sup>) | 25.64 | 0.88 | 0.0004 |  |  |  |
|  | Fe (ppm) | Cbrown (g/m<sup>2</sup>) |  |  |  | -1686.1 | 0.43 | 0.046 |
|  | Zn (ppm) | Cm (g/m<sup>2</sup>) |  |  |  | 0.05 | 0.41 | 0.051 |

Cbrown = Brown pigments content per area (g/m<sup>2</sup>); Car = Carotene content per area (g/m<sup>2</sup>); Cm = dry mass content per area. The results for the remaining (showing non-significant correlations) soil traits are in tables S1 and S2.

## Discussion

In this study, we set out to investigate (1) to what extent the spectral signature of leaves changes after removal from the tree and desiccation, (2) whether PROSPECT-D RTM tracks expected changes in leaf traits after leaf desiccation, (3) the influence of leaf state (fresh vs. dry) on PROSPECT-D parameter uncertainty, and (4) whether these spectrally predicted leaf traits can predict the variation of soil essential nutrients and chemical characteristics, and whether the state of the leaf (fresh vs. dry) impact the predictive capacity of these linear models. We expected the wavebands with highest degree of change within the spectral signature of dry leaves, relative to that of fresh leaves, to differ from Schweiger et al.’s (2018) local maxima. This was expected due to the displacement effect, expected under Wien’s law (Wien, 1897), which predicts a change of the waveband with the highest reflectance when the internal conditions of the leaf change (Jacquemoud & Ustin, 2019). The results showed that the maximum change in the visible spectrum (400 - 700 nm) and red edge (approximately at 720 nm) was sensed by shifts in the first derivative (slope) and second derivative (curvature) of the spectral signature, and at bands used for the prediction of photosynthetic pigments and lignin content (approximately at 400 nm and 720 nm). The maximum changes within the shortwave-infrared were sensed by changes in the reflectance intensity, as well as the slope and curvature of the spectral signature, in bands used for the prediction of protein content (∼1393 nm) and phenolic compounds (∼1400 nm), other bands related to foliar water content (∼1874) and the overtone of O-H bonds (∼1888 nm), which are related to water as well (Cheng et al., 2012; Couture et al., 2016; Curran, 1989; Fourty et al., 1996; Jin et al., 2017; Serbin & Townsend, 2020; S. Wang et al., 2020). These shifts observed in the spectral reflectance of desiccated leaves could influence trait predictability because of the covariation among leaf traits and the spectral reflectance that aids the ability of spectroscopy to predict leaf traits and, at the same time, worsen the prediction of others (Couture et al., 2016). Thus, the results show that by eliminating water and its disruptive effect, the spectral reflectance of leaves showed changes in other areas of the spectrum rather than at the local maxima for fresh leaves.

We asked if these changes have consequences for prediction precision of leaf traits based on leaf radiative transfer models. They do. The Bayesian inversion analysis of the PROSPECT model on fresh and desiccated leaves showed that this method is better at determining the leaf dry mass and brown pigments per area when leaves are dry. These results are not surprising since the components that PROSPECT defines as leaf dry mass (such as cellulose, starch and sugar content; Ustin & Jacquemoud, 2020) interact with the electromagnetic spectrum at the wavebands where water and proteins also interact (Carter, 1991; S. Wang et al., 2020). Therefore, it makes sense that the elimination of water and denaturation of proteins contribute to a better performance of the model. In the case of brown pigments, the accuracy of their content determination may be explained by their concentration - since these pigments emerge from protein decay and zinc accumulation (Féret et al., 2017). Regardless of the species, brown pigment accumulation within the leaf is a general trajectory in leaf senescence (Ustin & Jacquemoud, 2020), and that is what the results showed after the leaves were desiccated (Figure 4F). The opposite happened with photosynthetic pigments such as carotene, and equivalent water thickness since their content decreased after the leaves were desiccated (Figures 4 C and E). The structural N parameter, which describes the leaf’s internal structure, and where a higher number indicates higher complexity with more intercellular air space (Boren et al., 2019), showed an overall increase in the dry samples and no change in the uncertainty around the predicted values relative to models built from fresh leaves. An increase in this parameter is not surprising after desiccation, since water loss will leave more air space due to reduced water content and shrinkage of dead cells.

The results for the regression analysis linking leaf trait variation with soil nutrients suggest that models built with traits predicted from desiccated leaves are better at capturing the variation of some soil properties, which may be related to decreased prediction uncertainties. Brown pigment content, which measures the content of brown polyphenolic substance resulting from enzyme decomposition (Féret et al., 2008), was a good predictor for zinc in models built with dry leaves, which relates to the fact that plants produce these substances when zinc is in excess (Rauser, 1973). Brown pigment content was also good at predicting soil pH, boron, and iron in the models built with dry leaves. It is known that iron plays a role in the biosynthesis of lignin (Hajiboland, 2012), along with boron (Li et al., 2018). Furthermore, the capacity of the model to predict variation in soil pH may be a consequence of plant nutrient decrease: when plant nutrients decrease in the leaves, roots lower the soil pH to free soil nutrients, by releasing hydrogen protons (Guerinot & Yi, 1994). Other models showed carotene content as a good predictor for soil phosphorus, whose concentration in the soil impacts the regulation of carotene content (Xu & Mou, 2016). We also found that leaf dry matter content is a good predictor of soil sulfur, and there is empirical evidence of covariations between soil sulfur and leaf dry matter in *Arabidopsis* (Abbey et al., 2002). The only statistically significant models built from fresh leaves found that carotene content predicted soil manganese, which is a nutrient involved in the synthesis of carotenoids (Hajiboland, 2012). It is important to keep in mind that the lack of statistical significance in models built with fresh leaves may be due to an error caused by the disruptive effect of water over the prediction accuracy of spectrally derived traits.

## Conclusion

This work shows how leaf spectral reflectance, a complex characteristic, is affected by desiccation. In addition, it demonstrates that the prediction of important leaf traits such as leaf dry mass per area, a functional trait related to the capacity of plants to acquire carbon (van de Wega et al., 2009), improves with leaf desiccation. It also shows that the prediction of ecosystem processes such as nutrient transfer between the leaf and the soil, defined as the prediction of soil nutrient values by leaf traits, can be more precise when the models use leaf traits predicted from dry samples - which is possibly explained by the elimination of the disruptive power of water within the leaf. In this study, this improvement was observed in predictions of the structural parameter N, leaf dry mass, and brown pigments. Future research can use this investigation as a baseline for field-based explorations of the prediction of belowground processes. Larger sample sizes will likely contribute to strengthening the correlations between leaf traits and soil nutrients detected here. By demonstrating the contribution of desiccated leaves to studies of ecosystem functioning, this research opens the possibility of collecting data from substantially more species and areas, without the need of transporting a spectroradiometer to the field. It also highlights the usefulness of sampling living plant collections, such as the one harbored at the Enid A. Haupt Conservatory of the New York Botanical Garden, which can host a diverse assemblage of tropical plants in environments similar to those in their native ranges.

## Acknowledgments

We would like specially to thank Marc Hachadourian, Christian Primeau, and all the staff of the Enid A. Haupt Conservatory at the New York Botanical Garden for allowing us to work there before and during the COVID19 pandemic. We would also like to thank the people at the Soil Lab at the University of Connecticut for their wonderful work with their soil samples and to Maria Fernanda Chiappero from Universidad Nacional de Córdoba, Argentina for her suggestions on soil sampling methods. Finally, we would like to thank Amy Berkov, Ana Carnaval, and Fabián Michelangeli for helping with the edition of this manuscript.

## Author contributions

NLQC, SPS, and DCD designed the study. NLQC and SPS performed field sampling. NQLC performed the statistical analyses and wrote the first draft of the manuscript. All authors contributed to the interpretation of results and revisions of the manuscript.

## Data availability

## Supplementary Information

**Table S1.**
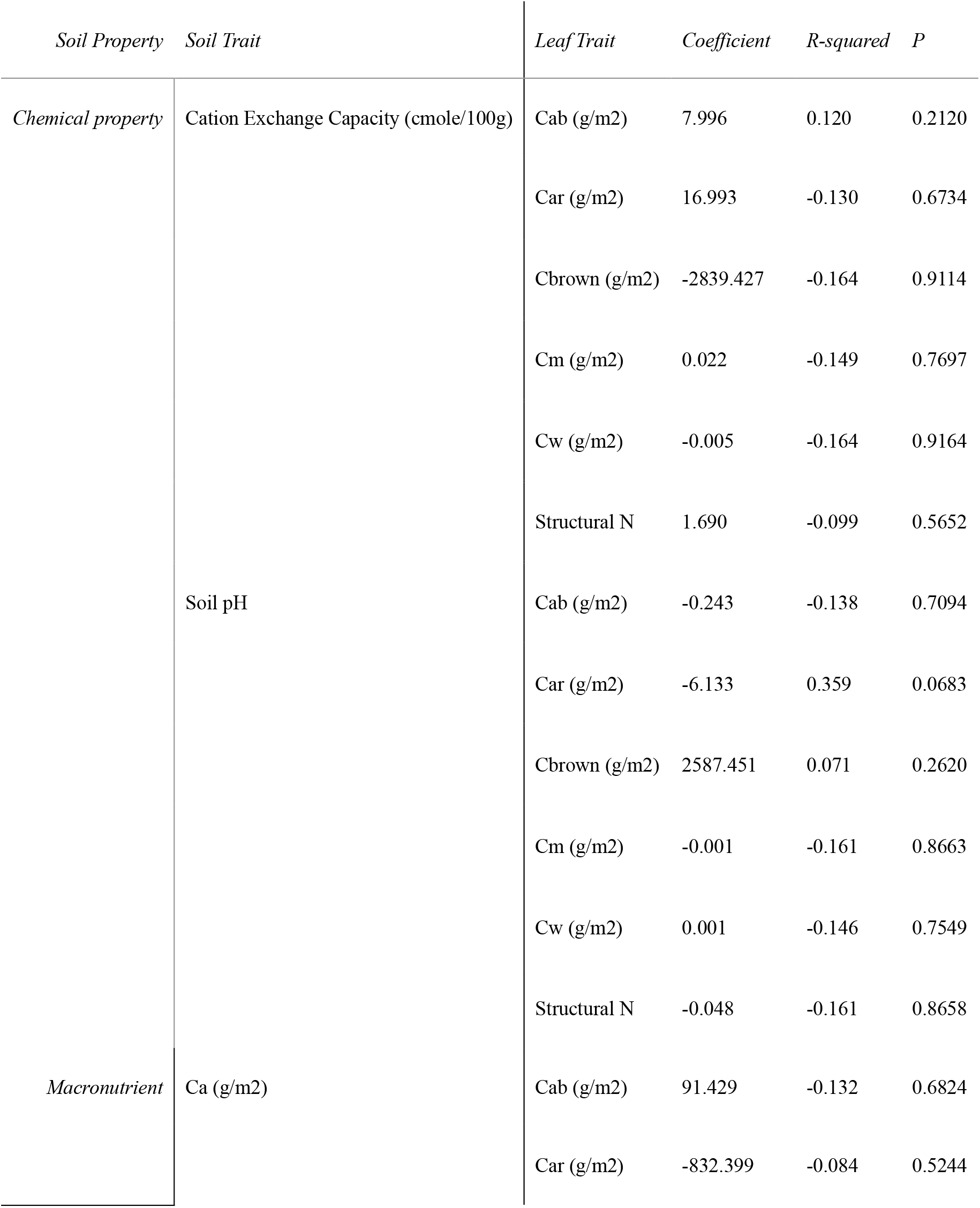
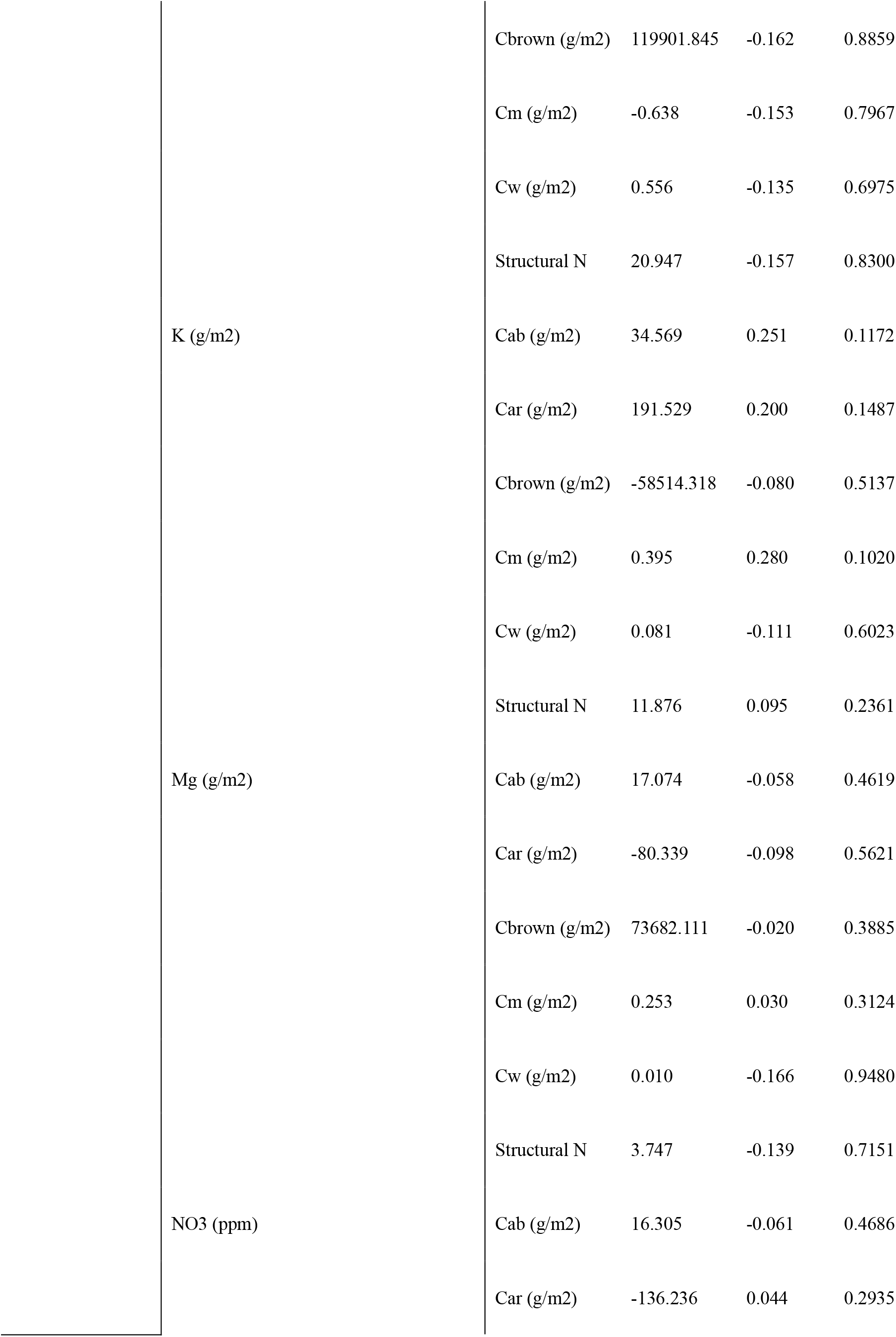
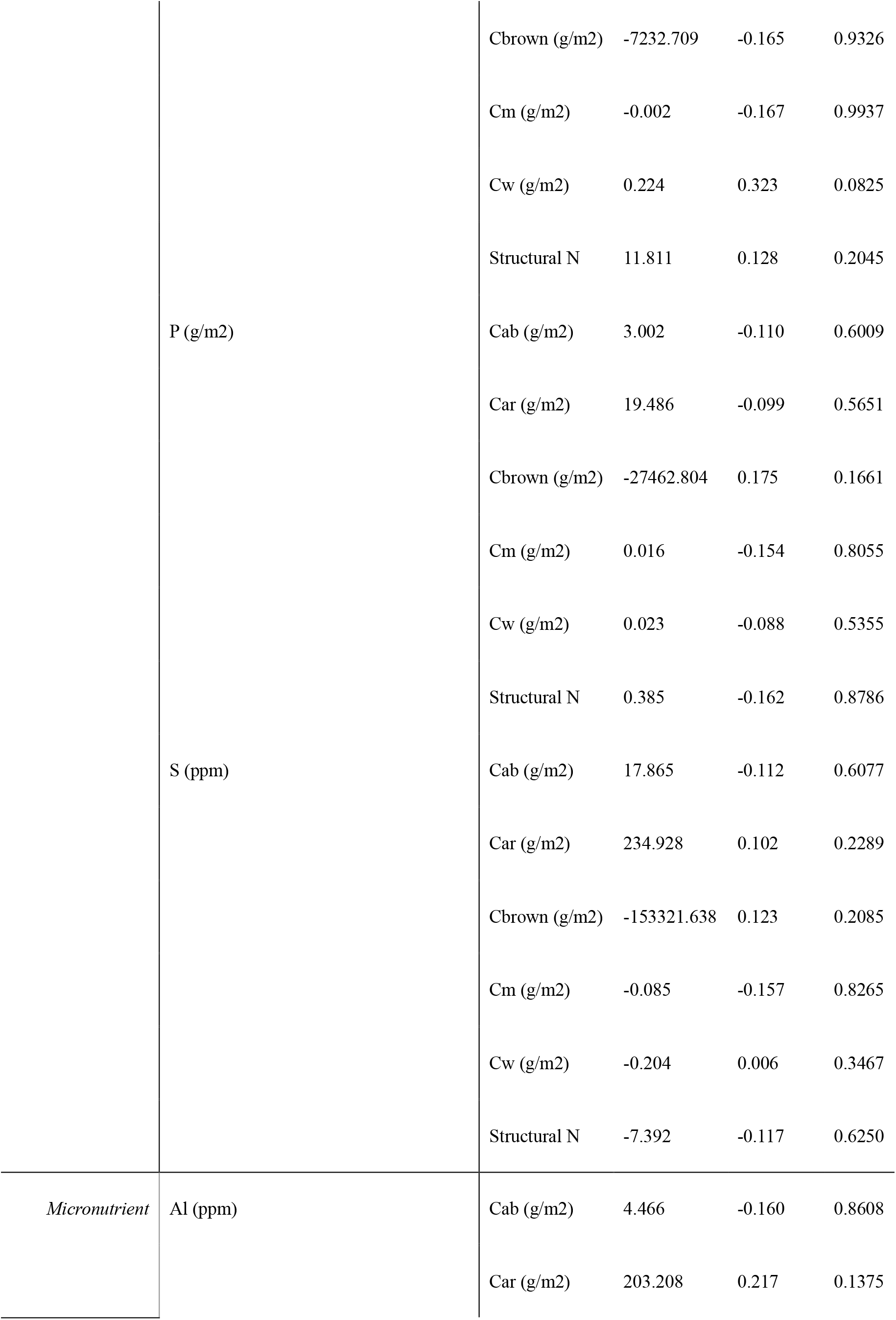
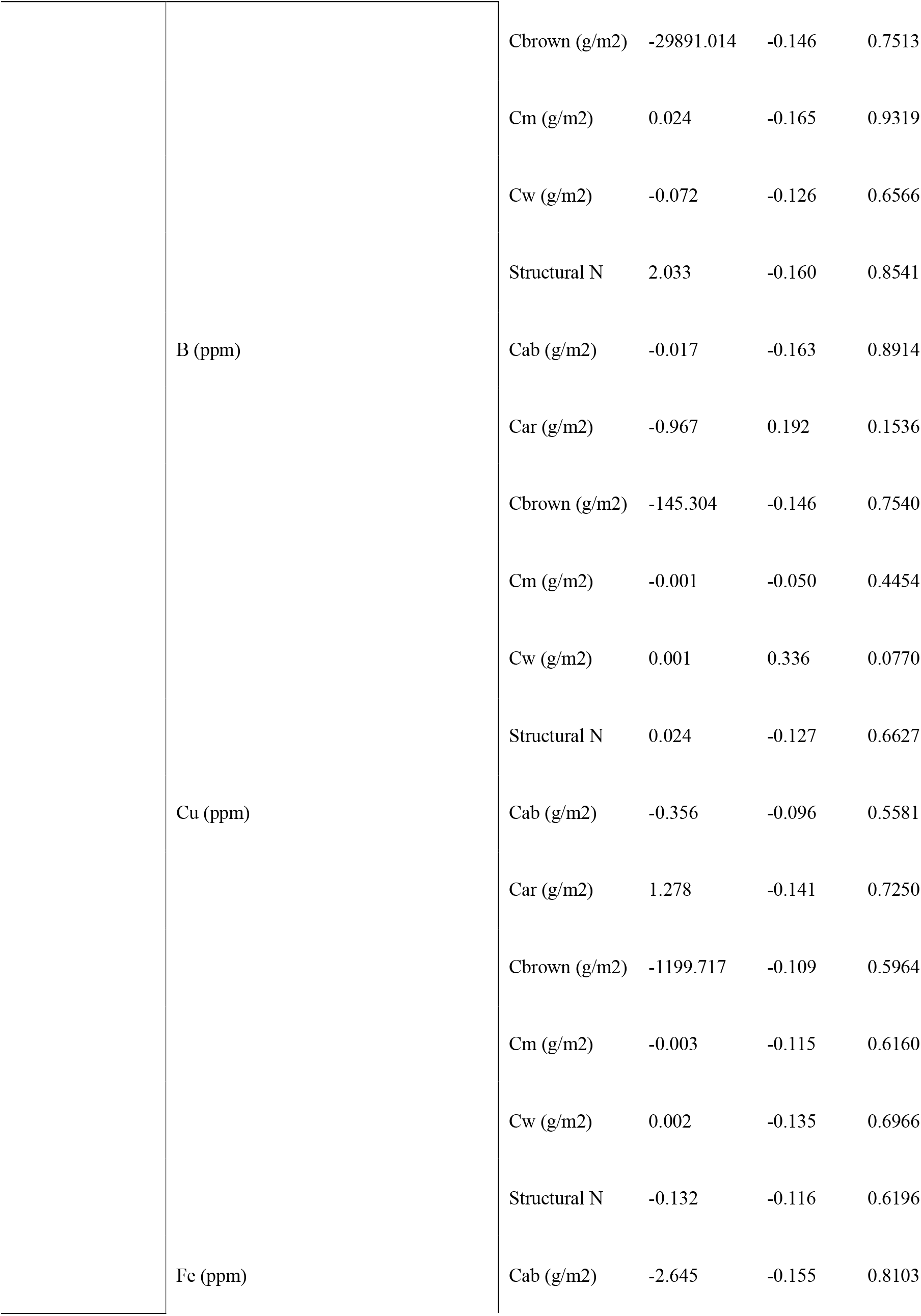
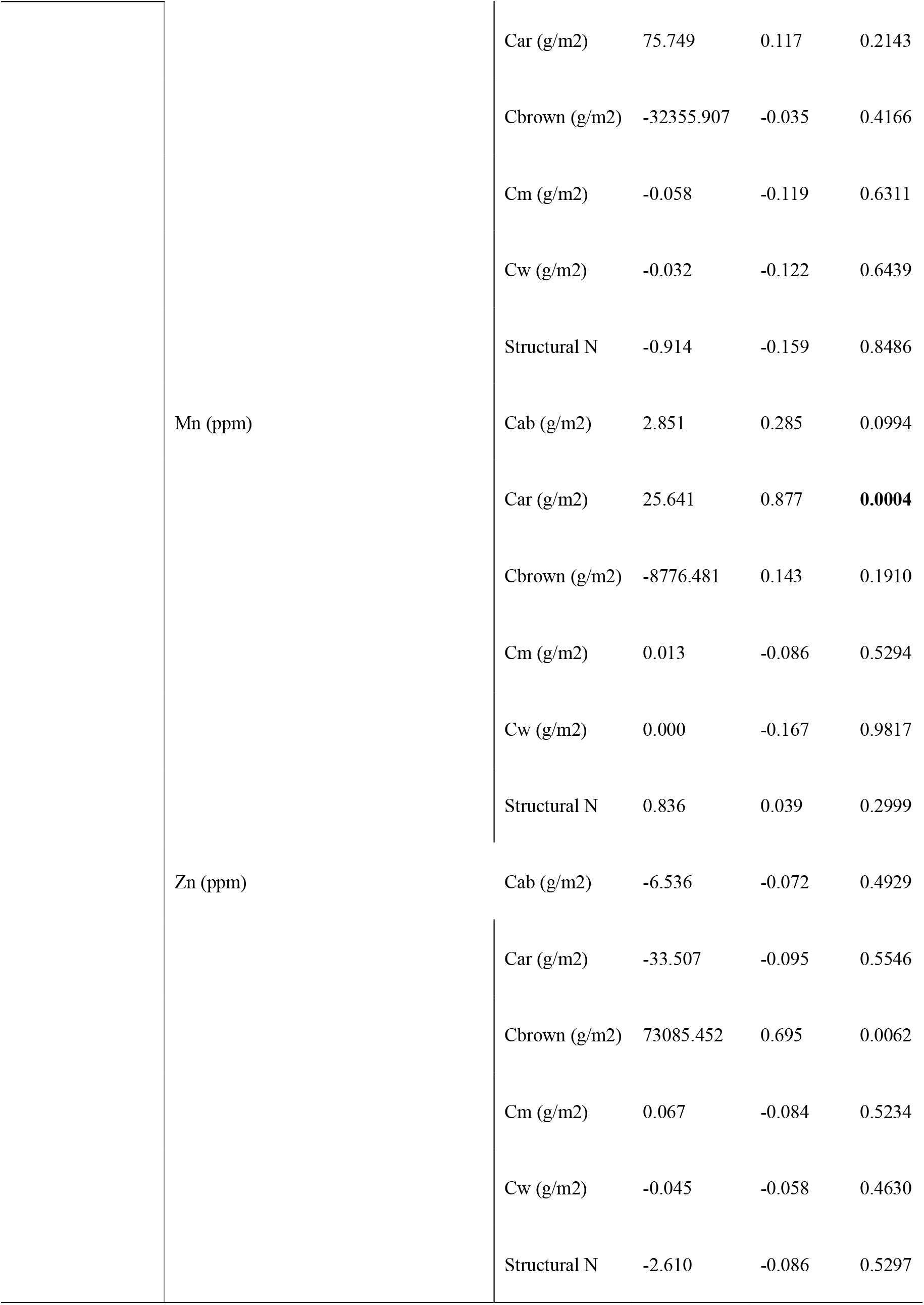
Soil – leaf relationships, based on a bivariate linear model, derived from fresh specimens. Cab = total chlorophylls content per area (g/m2); Cbrown = Brown pigments content per area (g/m2); Car = Carotene content per area (g/m2); Cm = dry mass content per area; Cw = equivalent water thickness (g/m2).

**Table S2.**
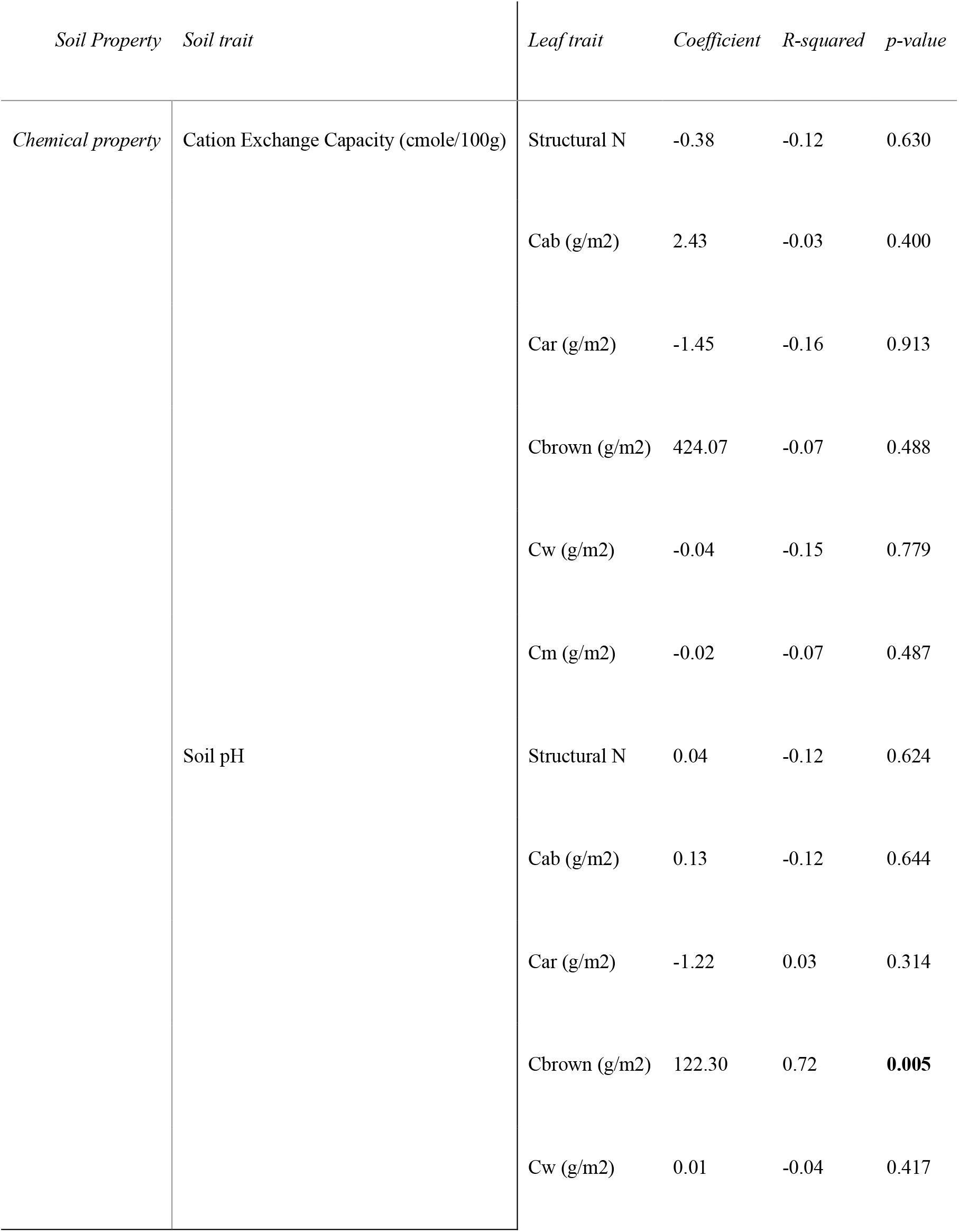
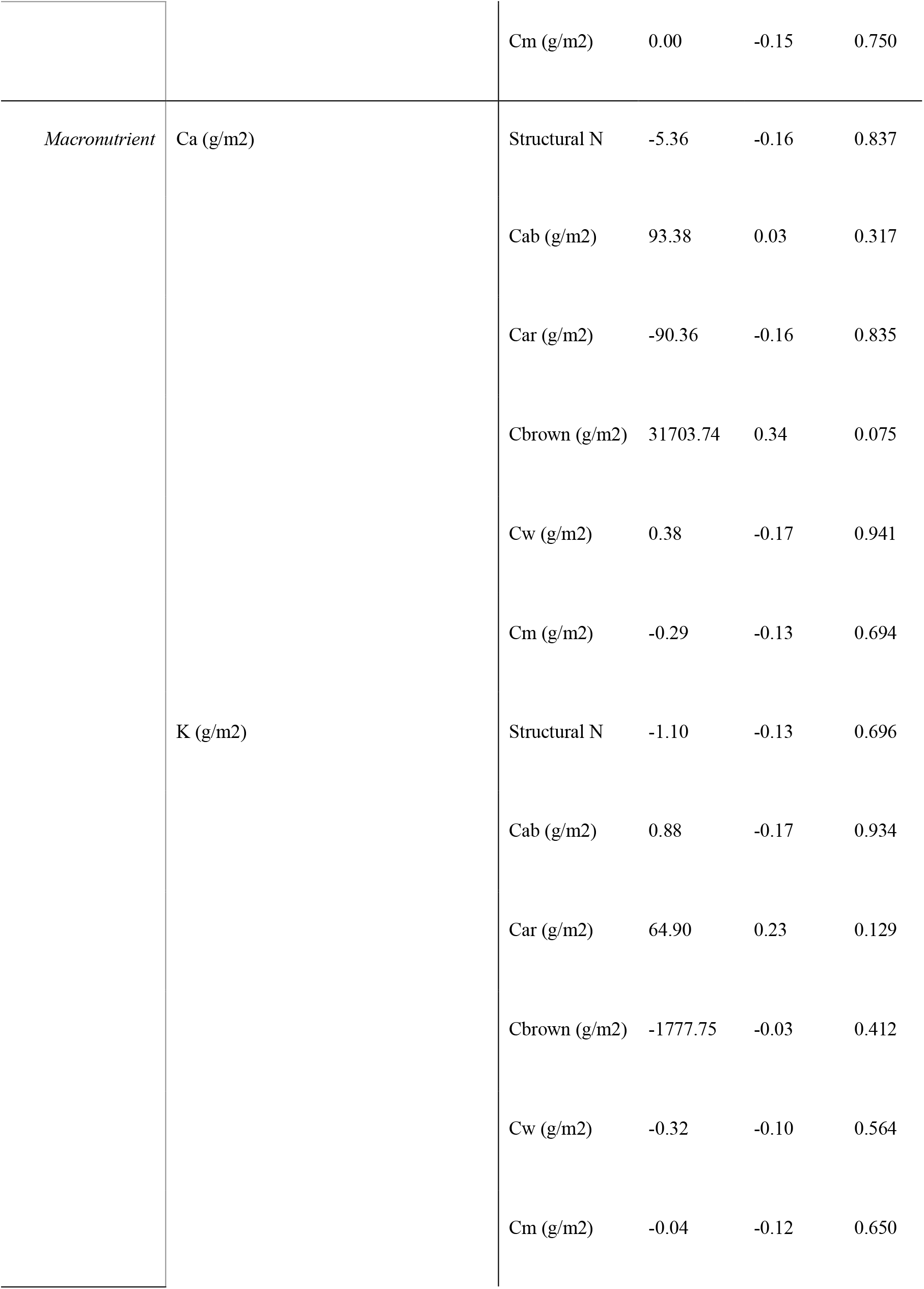
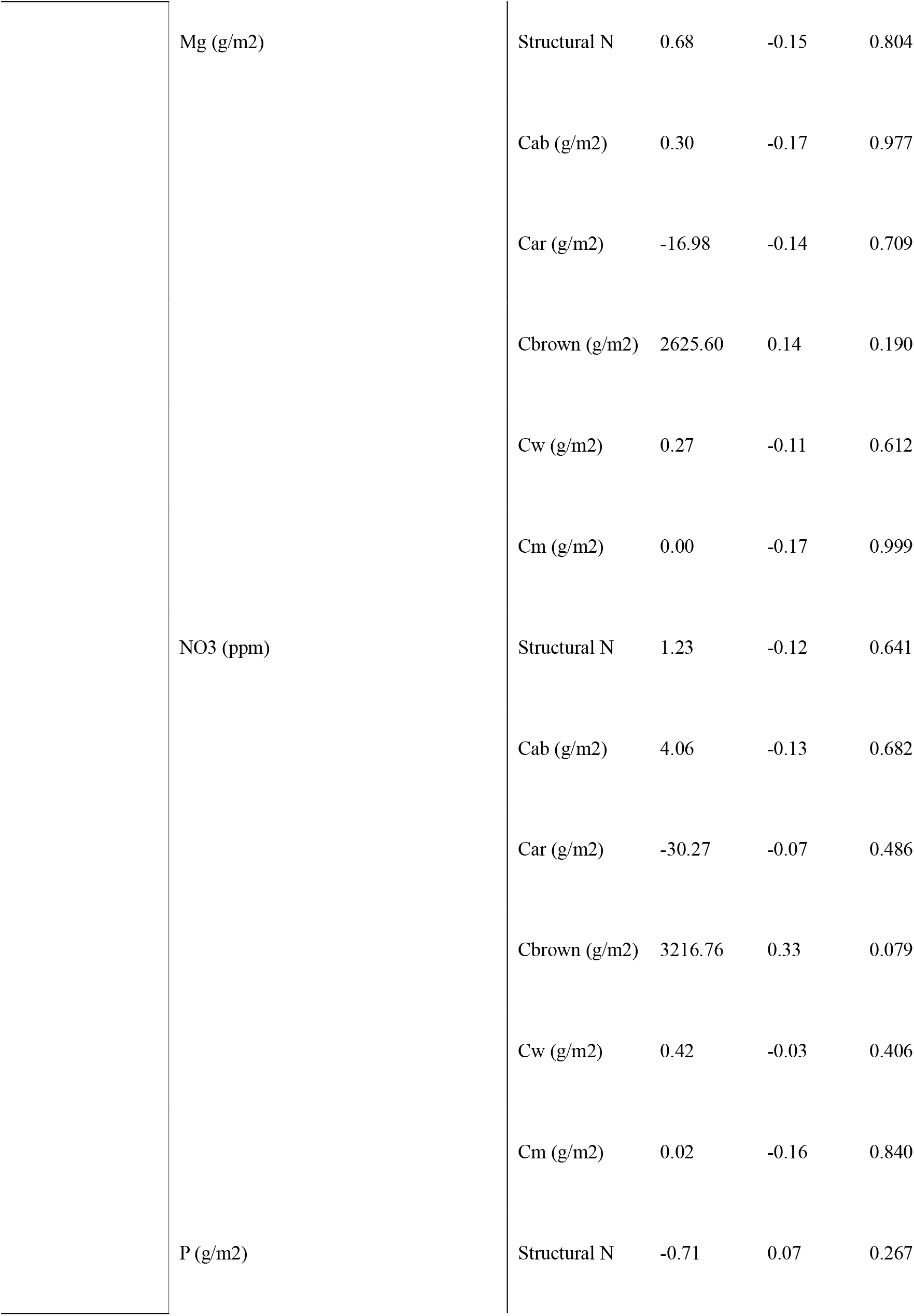
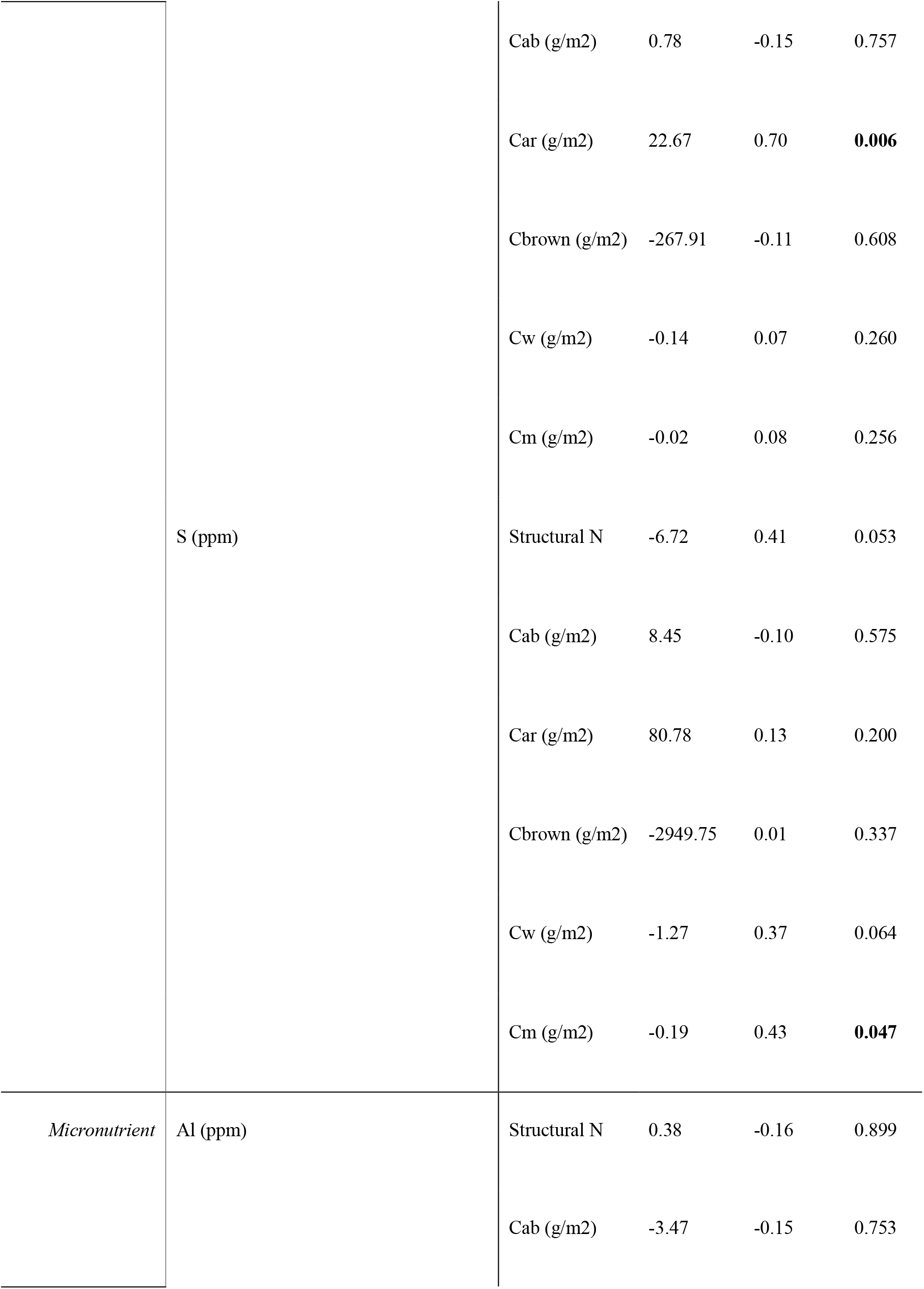
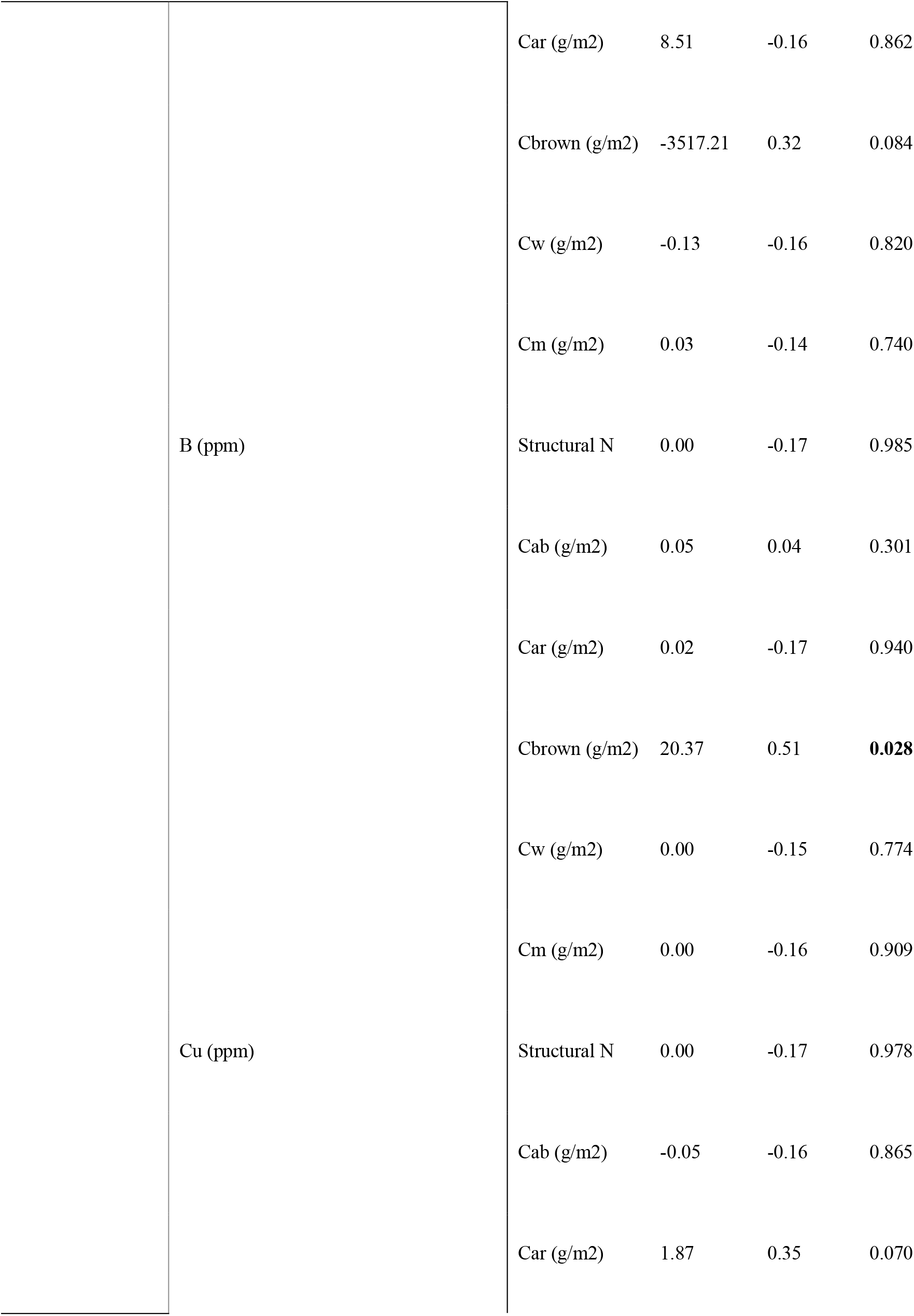
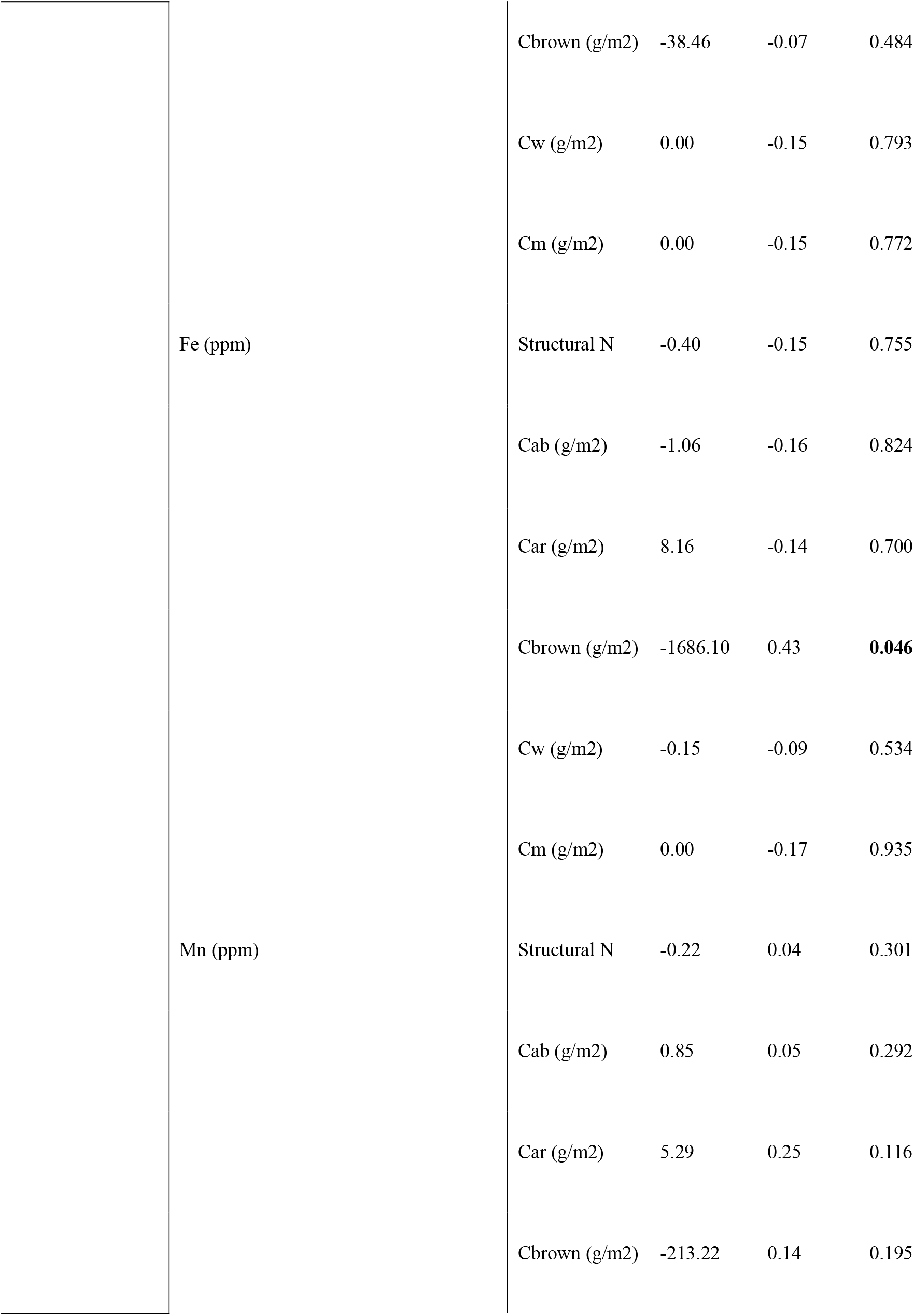
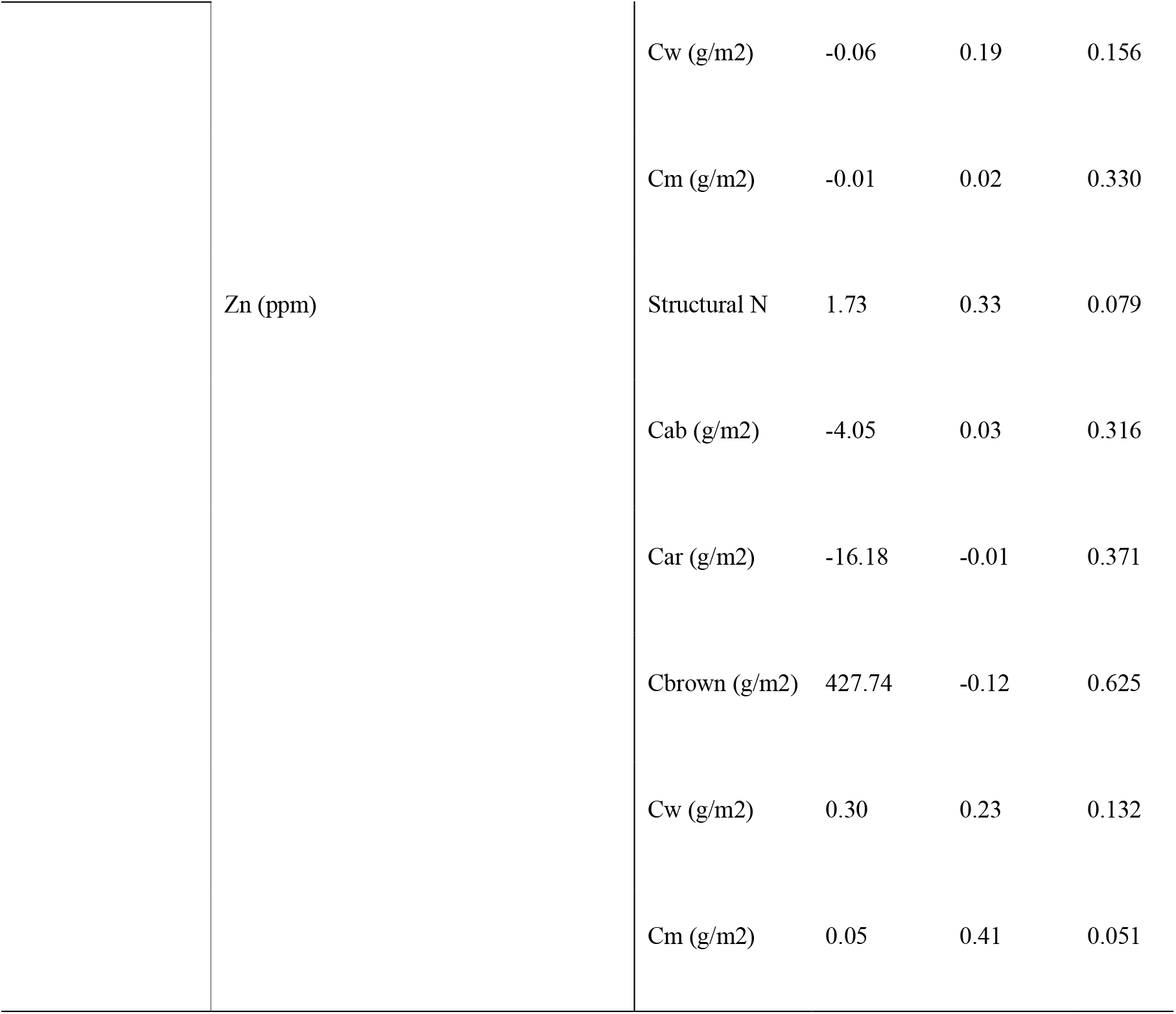
Soil – leaf relationships, based on a bivariate linear model, derived from desiccated specimens where Cab = Total chlorophylls content per area (g/m^2^); Cbrown = Brown pigments content per area (g/m^2^); Car = Carotene content per area (g/m^2^); Cm = dry mass content per area; Cw = equivalent water thickness (g/m^2^).

**Figure S1.**
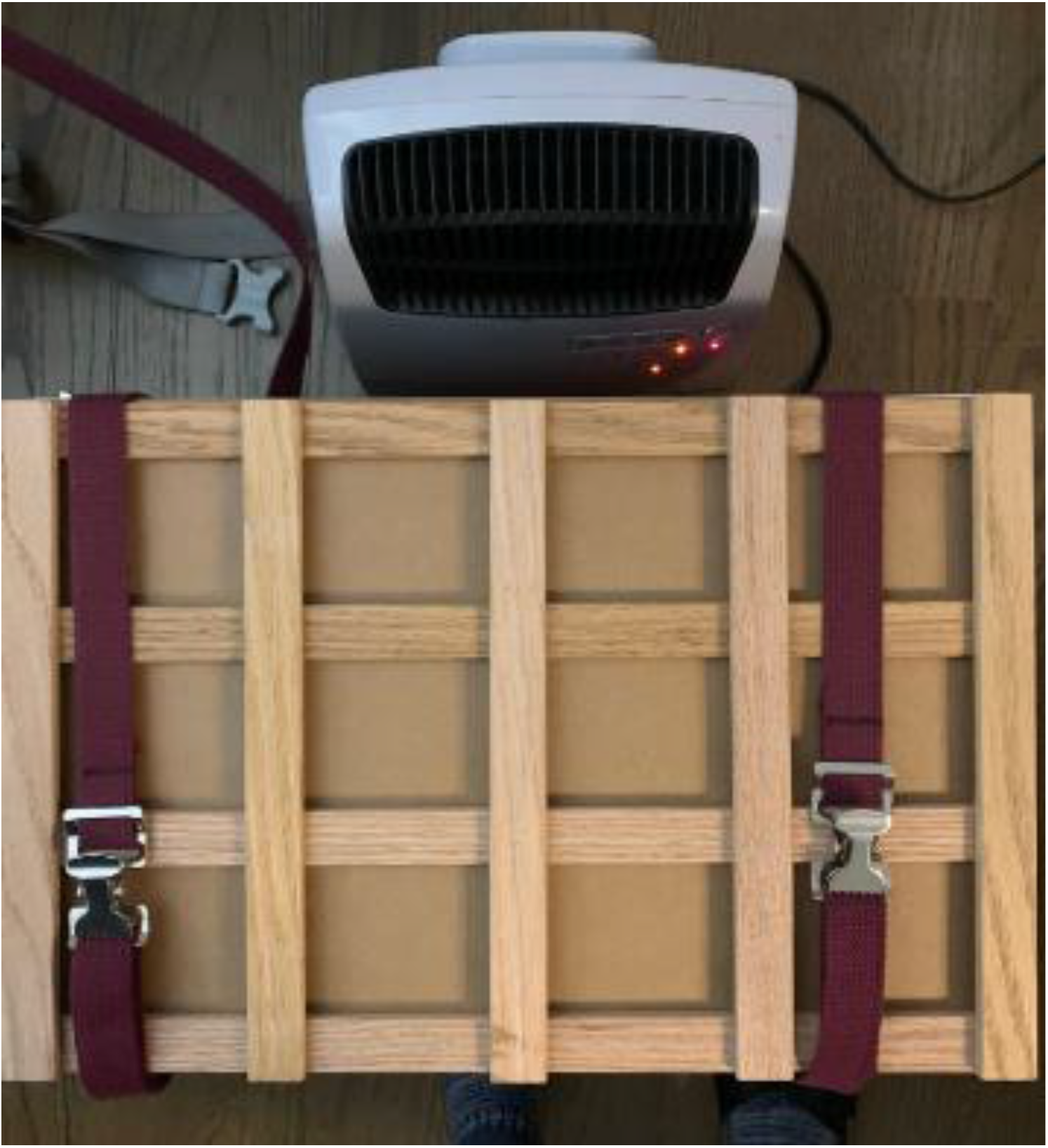
Leaf drying system implemented in this study. Leaves were placed on C-flute corrugated ventilators and pressed with a botanical field press while placed in front of a small Lasko® space heater.

**Figure S2.**
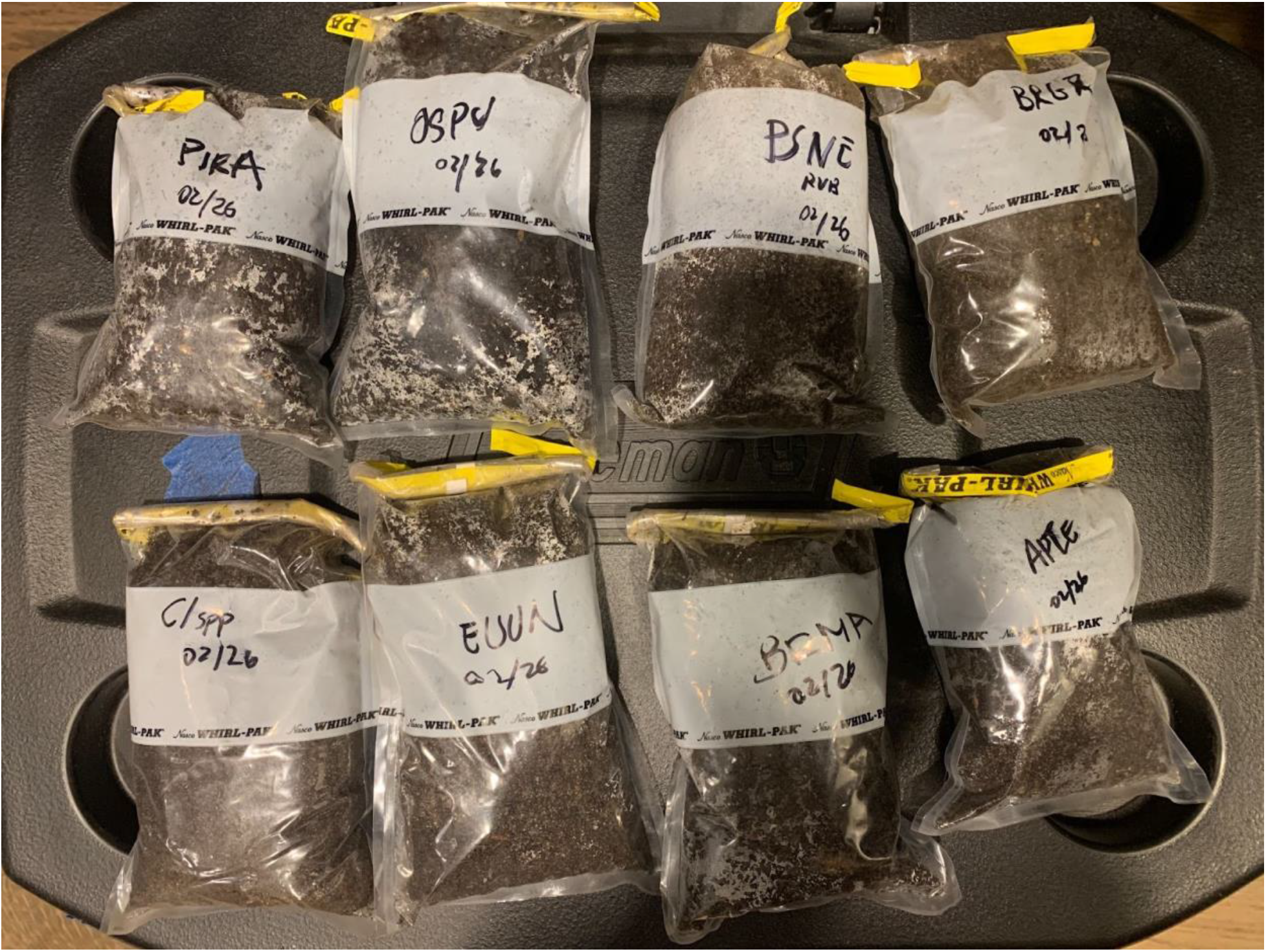
Soil samples from rhizosphere for *Osa pulchra* (OSPU), *Pimenta racemosa* (PIRA), *Psychotria nervosa* (PSNE), *Brunfelsia grandiflora* (BRGR), *Cinchona spp* (CIspp), *Eugenia uniflora* (EUUN), *Brownea macrophylla* (BRMA), *Aphelandra tetragona* (APTE).

